# Dynamic interaction network inference from longitudinal microbiome data

**DOI:** 10.1101/430462

**Authors:** Jose Lugo-Martinez, Daniel Ruiz-Perez, Giri Narasimhan, Ziv Bar-Joseph

## Abstract

**Background:** Several studies have focused on the microbiota living in environmental niches including human body sites. In many of these studies researchers collect longitudinal data with the goal of understanding not just the composition of the microbiome but also the interactions between the different taxa. However, analysis of such data is challenging and very few methods have been developed to reconstruct dynamic models from time series microbiome data.

**Results:** Here we present a computational pipeline that enables the integration of data across individuals for the reconstruction of such models. Our pipeline starts by aligning the data collected for all individuals. The aligned profiles are then used to learn a dynamic Bayesian network which represents causal relationships between taxa and clinical variables. Testing our methods on three longitudinal microbiome data sets we show that our pipeline improve upon prior methods developed for this task. We also discuss the biological insights provided by the models which include several known and novel interactions.

**Conclusions:** We propose a computational pipeline for analyzing longitudinal microbiome data. Our results provide evidence that microbiome alignments coupled with dynamic Bayesian networks improve predictive performance over previous methods and enhance our ability to infer biological relationships within the microbiome and between taxa and clinical factors.

## 1. Introduction

Multiple efforts have attempted to study the microbiota living in environmental niches including human body sites. These microbial communities can play beneficial as well as harmful roles in their hosts and environments. For instance, microbes living in the human gut perform numerous vital functions for homeostasis ranging from harvesting essential nutrients to regulating and maintaining the immune system. Alternatively, a compositional imbalance known as dysbiosis can lead to a wide range of human diseases [1], and is linked to environmental problems such as harmful algal blooms [2].

While many studies profile several different types of microbial taxa, it is not easy in most cases to uncover the complex interactions within the microbiome and between taxa and clinical factors (e.g., gender, age, ethnicity). Microbiomes are inherently dynamic, thus, in order to fully reconstruct these interactions we need to obtain and analyze longitudinal data [3]. Examples include characterizing temporal variation of the gut microbial communities from pre-term infants during the first weeks of life, and understanding responses of the vaginal microbiota to biological events such as menses. Even when such longitudinal data is collected, the ability to extract an accurate set of interactions from the data is still a major challenge.

To address this challenge we need computational time-series tools that can handle data sets that may exhibit missing or noisy data and non-uniform sampling. Furthermore, a critical issue which naturally arises when dealing with longitudinal biological data is that of temporal rate variations. Given longitudinal samples from different individuals (for example, gut microbiome), we cannot expect that the rates in which interactions take place is exactly the same between these individuals. Issues including age, gender, external exposure, etc. may lead to faster or slower rates of change between individuals. Thus, to analyze longitudinal data across individuals we need to first align the microbial data. Using the aligned profiles we can next employ other methods to construct a model for the process being studied.

Most current approaches for analyzing longitudinal microbiome data focus on changes in outcomes over time [4, 5]. The main drawback of this approach is that individual microbiome entities are treated as independent outcomes, hence, potential relationships between these entities are ignored. An alternative approach involves the use dynamical systems such as the generalized Lotka-Volterra (gLV) models [6, 7, 8, 9]. While gLV and other dynamical systems can help in studying the stability of temporal bacterial communities, they are not well-suited for causality and probabilistic inference over discrete time. Finally, probabilistic graphical models (e.g., hidden Markov models, Kalman filters and dynamic Bayesian networks) are machine learning tools which can effectively model dynamic processes, as well as discover causal interactions [10].

In this work we first adapt statistical spline estimation and dynamic time-warping techniques for aligning time-series microbial data so that they can be integrated across individuals. We use the aligned data to learn a Dynamic Bayesian Network (DBN), where nodes represent microbial taxa, clinical conditions, or demographic factors and edges represent causal relationships between these entities. We evaluate our model by using multiple data sets comprised of the microbiota living in human body parts including gastrointestinal tract, urogenital tract and oral cavity. We show that models for these systems can accurately predict changes in taxa and that they greatly improve upon models constructed by prior methods. Finally, we characterize the biological relationships in the reconstructed microbial communities and discuss known and novel interactions discovered by these models.

## 2. Methods

### Data sets

We collected multiple public longitudinal microbiome data sets for testing our method:

#### Infant gut microbiome

This data set was collected by La Rosa *et al*. [5]. They sequenced gut microbiome from 58 pre-term infants in neonatal intensive care unit (NICU). The data was collected during the first 12 weeks of life (until discharged from NICU or deceased) sampled every day or two on average. Following analysis 29 microbial taxa were reported across the 922 total infant gut microbiome measurements. In addition to the taxa information, this data set includes clinical and demographic information for example, gestational age at birth, post-conceptional age when sample was obtained, mode of delivery (C-section or vaginal), antibiotic use (percentage of days of life on antibiotic), and more (see Additional file 1: Table S1 for complete list of clinical features available).

#### Vaginal microbiome

The vaginal microbiota data set was collected by Gajer *et al*. [4]. They studied 32 reproductive-age healthy women over a 16-week period. This longitudinal data set is comprised of 937 self-collected vaginal swabs and vaginal smears sampled two times a week. Analysis identified 330 bacterial taxa in the samples. The data also contains clinical and demographic attributes on the non-pregnant women such as Nugent score [11], menses duration, tampon usage, vaginal douching, sexual activity, race and age. To test the alignment methods we further sub-divided the microbial composition profiles of each subject by menstrual periods. This resulted in 119 time-series samples, an average of 3–4 menstrual cycles per woman. Additional file 2: Figure S1a shows four sub-samples derived from an individual sample over the 16-week period along with corresponding menses information.

#### Oral cavity microbiome

The cavity data was downloaded from the case-control study conducted by DiGiulio *et al*. [12] comprised of 40 pregnant women, 11 of whom delivered pre-term. Overall they collected 3, 767 samples and identified a total of 1, 420 microbial taxa. Data was collected weekly during gestation and monthly after delivery from four body sites: vagina, distal gut, saliva, and tooth/gum. In addition to bacterial taxonomic composition, these data sets report clinical and demographic attributes which include gestational status, gestational or postpartum day when sample was collected, race and ethnicity. In this paper, we solely focus on the tooth/gum samples during gestation from Caucasian women in the control group to reduce potential confounding factors. This restricted set contains 374 temporal samples from 18 pregnant women.

Additional file 1: Table S1 summarizes the three longitudinal microbiome data sets used in this study, including the complete list of clinical features available.

### Temporal alignment

As mentioned in the Background, a challenge when comparing time series obtained from different individuals is the fact that while the overall process studied in these individuals may be similar, the *rates* of change may differ based on several factors (age, gender, other diseases, etc.). Thus, prior to modeling the relationships between the different taxa we first align the data sets between individuals by warping the time scale of each sample into the scale of another representative sample referred to as *reference*. The goal of an alignment algorithm is to determine, for each individual *i*, a function *f_i_(t)* which takes as an input a reference time *t* and outputs the corresponding time for individual *i*. Using this function we can compare taxa values for all individuals sampled for the same time point. This approach effectively sets the stage for accurate discovery of trends and patterns, hence, further disentangling the dynamic and temporal relationships between entities in the microbiome.

There are several possible options for selecting transformation function *f_i_*. Most methods used to date rely on polynomial functions [13, 14]. Prior work on the analysis of gene expression data indicated that given the relatively small number of time points for each individual simpler functions tend to outperform more complicated ones [15]. Therefore, we used a first degree polynomial: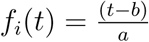 as the alignment function for tackling the temporal alignment problem, where *a* and *b* are the parameters of the function.

### Data pre-processing

Since alignment relies on continuous (polynomial) functions while the data is sampled at discrete internals, the first step is to represent the sample data using continuous curves as shown by the transition from Fig. 1a to Fig. 1b. Following prior work [15], we use B-splines for fitting continuous curves to microbial composition time-series data, thus, enabling principled estimation of unobserved time points and interpolation at uniform intervals. To avoid overfitting we removed any sample that had less than nine measured time points, and estimated a cubic B-spline from the observed abundance profile for all taxa in remaining samples using *splrep* and *BSpline* from the Python function *scipy.interpolate*. Additional file 3: Figure S2 shows the original and cubic spline of a representative microbial taxa from a randomly selected individual sample across each data set.

**Figure 1:**
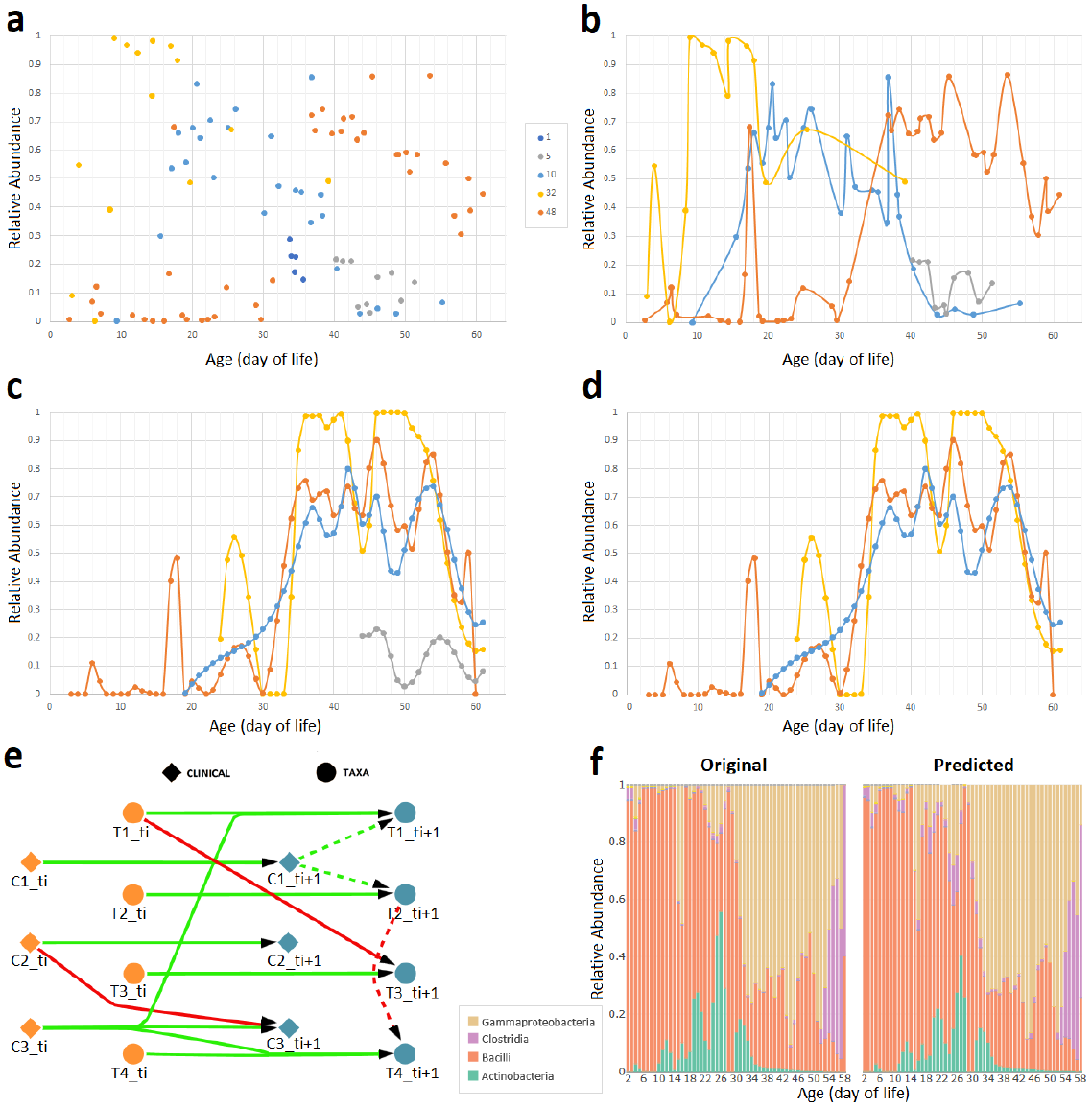
Computational pipeline proposed in this work. Figure shows microbial taxa Gammaproteobacteria at each step in the pipeline from a set of five representative individual samples (subjects 1, 5, 10, 32 and 48) of the infant gut data set at a sampling rate of 1 day**. a** — Raw relative abundance values for each sample measured at (potentially) non-uniform intervals even within the same subject**. b** — Cubic B-spline curve for each individual sample. Sample corresponding to subject 1 (dark blue) contains less than predefined threshold for measured time points, thus, removed from further analysis. The remaining smoothed curves enable principled estimation of unobserved time points and interpolation at uniform intervals**. c** — Temporal alignment of each individual sample against a selected reference sample (subject 48 shown in orange)**. d** — Post-alignment filtering of samples with alignment error higher than a pre-defined threshold. Sample corresponding to subject 5 (grey) discarded**. e** — Learning a dynamic Bayesian network (DBN) structure and parameters. Let nodes (*T*_1_,*T*_2_,*T*_3_,*T*_4_) represent microbial taxa and (*C*_1_, *C*_2_, *C*_3_) represent clinical factors shown as circles and diamonds, respectively. Figure shows two consecutive time slices *ti* and *ti* + 1, where dotted lines connect nodes from the same time slice referred to as intra edges, and solid lines connect nodes between time slices referred to as inter edges. Biological relationships are inferred from edge parameters in the learned DBN which can be positive (green) or negative (red)**. f** — Original and predicted relative abundance across four infant gut taxa for subject 48 at sampling rate of 1 day. Performance is evaluated by average mean absolute error (MAE) between original and predicted abundance values (MAE = 0.011).

### Aligning microbial taxon

To discuss the alignment algorithm we first assume that a reference sample, to which all other samples would be aligned, is available. We next discuss how to chose such reference.

Formally, let 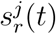 be the spline curve for microbial taxa *j* at time 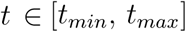 in the reference time-series sample *r*, where *t_min_* and *t_max_* denote the starting and ending time points of 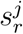, respectively. Similarly, let 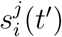 be the spline for individual *i* in the set of samples to be warped for taxa *j* at time 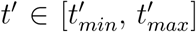. Next, analogously to Bar-Joseph *et al*. [13], the alignment error for microbial taxa *j* between 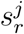 and 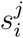 is defined as

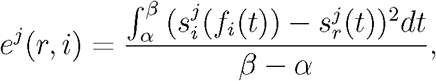

where 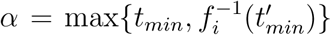 and 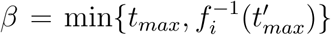 correspond to the starting and ending time points of the alignment. Observe that by smoothing the curves, it is possible to estimate the values at any intermediate time point in the alignment interval [*α*, *β*]. Finally, we define the microbiome alignment error for a microbial taxon of interest *S* between individual samples *r* and *i* as follows

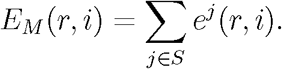

Given a reference *r* and microbial taxon *S*, the alignment algorithm task is to find parameters *a* and *b* that minimize *E_M_* for each individual sample *i* in the data set subject to the constraints: 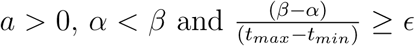. The latter constraint enforces that the overlap between aligned interval [*a*, *β*] and reference interval [*t_min_*, *t_max_]* is at least *ϵ*, otherwise trivial solutions (for example, no overlap leading to 0 error) would be selected. Here we used *ϵ* = 0.3 though results remain the same with larger values of *ϵ*. Fig. 1c illustrates an aligned set of four samples where reference sample *r* is shown in orange. Alternatively, Additional file 2: Figure S1b shows the temporal alignment between the sub-samples of the vaginal microbiome sample shown in Figure S1a for taxa *L. crispatus* using the first menstrual period sub-sample as reference (shown in orange).

### Selecting a reference sample

Finding a reference that jointly minimizes *E_m_* for all samples requires combinatorial analysis which takes time that is exponential in the number of individuals [13].

Instead, we used a heuristic approach to find the best pairwise alignment function and, ultimately, select an optimal reference. In particular, we first find the best pairwise alignments via a grid-search parameter sweep between *a* ∈ (0, 4] with increments of 0.01 and *b* ∈ [–50, 50] with increments of 0.5 in the linear alignment function *f_i_* previously described. It is important to note that this restricted search space for parameters *a* and *b* may lead to some sample pairs (*r*, *i*) without a temporal alignment because overlap constraint is not met. Additionally, we filtered out any microbial taxa *j* ∈ *S* for which the mean abundance in either 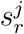 or 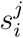 was less than 0.1%, or had zero variance over the originally sampled time points. Lastly, an optimal reference for each data set is determined by generating all possible pairwise alignments between samples. To select the best reference *r*\* we employed the following criteria: (1) at least 90% of the individual samples are aligned to *r\**, and (2) the alignment error *E_M_* is minimized.

#### Abnormal or noisy samples filtering

As a post-processing step, we implemented a simple procedure which takes as input the resulting individual-wise alignments to identify and filter out abnormal and noisy samples. Given an aligned microbiome data set, we (1) computed the mean *μ* and standard deviation *δ* of the alignment error *E_M_* across all aligned individual samples, and (2) removed all samples from an individual where 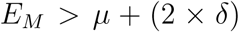. Fig. 1d shows the filtered set for the aligned taxa in the previous step (Fig. 1c). This analysis can both, help to identify outliers and improves the ability to accurately reconstruct models for taxa interactions as we show in Results.

#### Taxon selection from alignment

As previously described, the microbiome alignment error *E_M_* for a pairwise alignment is restricted to the set of microbial taxa *S* which contributed to the alignment. However, this set of microbes can vary for different pairwise alignments even with the same reference. Therefore, we focused on the subset of taxa which contributed to at least half of the pairwise alignments for the selected reference.

Additional file 4: Table S2 lists alignment information for each data set such as reference sample, number of aligned samples and selected taxa.

### Model construction

Using the aligned taxa, we next attempted to learn graphical models that provide information about the causal impacts of taxa and clinical or demographic variables on other taxa. For this, we used Dynamic Bayesian Networks (DBNs) which have been widely used to model sequential data, including speech [16, 17], biological [18, 19, 10], or economic sequences [20, 21]. A DBN is a directed acyclic graph where, at each *time slice* (or time instance), nodes correspond to random variables of interest (e.g., taxa, post-conceptional age, or Nugent score) and directed edges correspond to their conditional dependencies in the graph [22]. These time slices are not modeled separately. Instead a DBN contains edges connecting time slices known as *inter edges* that are repeated for each time point modeled as depicted in Fig. 1e. In summary, the model learns the transition probability from one time point to the next as a stationary conditional probability. DBNs are considered generative models, therefore, ideal for modeling the compositional interactions and dynamics of the microbiota given the first time point.

Here, we use a “two-stage” DBN model in which only two slices are modeled and learned at a time. Throughout this paper, we will refer to the previous and current time slice with suffix *ti* and *ti* + 1, respectively. Fig. 1e illustrates a skeleton of the general structure of a two-stage DBN in the context of a longitudinal microbiome study. In this example, for each time slice, the nodes correspond to random variables of observed quantities for different microbial taxa (*T*_1_, *T*_2_, *T*_3_, *T*_4_) or clinical factors (*C*_1_, *C*_2_, *C*_3_) shown as circles and diamonds, respectively. These variables can be connected by intra edges (dotted lines) or inter edges (solid lines). In this DBN model, the abundance of a particular microbe in the current time slice is determined by parameters from both intra and inter edges, thus, modeling the complex interactions and dynamics between the entities in the microbial community.

Typically, analysis using DBNs is divided into two components: learning the network structure and parameters and inference on the network. The former can be further sub-divided into (i) structure learning which involves inferring from data the causal connections between nodes (i.e., learning the intra and inter edges) while avoiding overfitting the model, and (ii) parameter learning which involves learning the parameters of each intra and inter edge in a specific network structure. There are only a limited number of open software packages which support both learning and inference with DBNs [23, 24] in the presence of discrete and continuous variables. Here we used CGBayesNets package [23, 10] which is freely available software package for learning the network structure and performing inference for Conditional Gaussian Bayesian models [25]. While useful, CGBayesNets does not support several aspects of DBN learning including the use of intra edges, searching for a parent candidate set in the absence of prior information and more. We have thus extended the structure learning capabilities of CGBayesNets to include intra edges while learning network structures and implemented well-known network scoring functions for penalizing models based on the number of parameters such as Akaike Information Criterion (AIC) and Bayesian Information Criterion (BIC) [26].

#### Learning DBN model parameters

Let Θ denote the set of parameters for the DBN and *G* denote a specific network structure over discrete and continuous variables in the microbiome study. In a similar manner to McGeachie *et al*. [10], we can decompose the joint distribution as

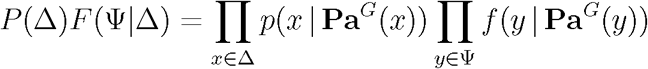

where *P* denotes a set of conditional probability distributions over discrete variables Δ, *F* denotes a set of linear Gaussian conditional densities over continuous variables Ѱ, and **Pa***^G^*(*X*) denotes the set of parents for variable *X* in *G*. Since we are dealing with both, continuous and discrete nodes in the DBN, in our method, continuous variables (i.e., microbial taxa compositions) are modeled using a Gaussian with the mean set based on a regression model over the set of continuous parents as follows

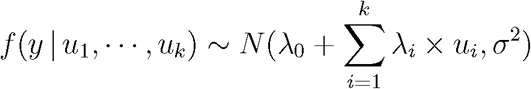

where *u_1_*, …, *u_k_* are continuous parents of *y*; *λ*_0_ is the intercept; *λ*_1_, … *,λ_k_* are the corresponding regression coefficients for *u_1_*, …, *u_k_*; and *σ*^2^ is the standard deviation. We point out that if *y* has discrete parents then we need to compute coefficients 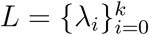 and standard deviation *σ*^2^ for each discrete parents configuration. For example, the conditional linear Gaussian density function for variable 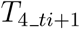 in Fig. 1e denoted as 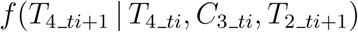 is modeled by

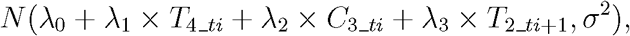

where *λ*_1_, *λ*_2_, *λ*_3_ and *σ^2^* are the DBN model parameters. In general, given a longitudinal data set *D* and known structure *G*, we can directly infer the parameters Θ by maximizing the likelihood of the data given our regression model.

#### Learning DBN structure

Learning the DBN structure can be expressed as finding the optimal structure and parameters

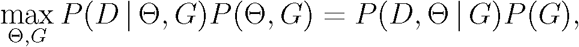

where *P*(*D* | Θ, *G*) is the likelihood of the data given the model. Intuitively, the likelihood increases as the number of valid parents **Pa***^G^*(·) increases, thus, making it challenging to infer the most accurate model for data set *D*. Therefore, the goal is to effectively search over possible structures while using a function that penalizes overly complicated structures and protects from overfitting.

Here, we maximize *P(D*, Θ | *G*) for a given structure *G* using maximum likelihood estimation (MLE) coupled with BIC score instead of Bayesian Dirichlet equivalent sample-size uniform (BDeu) metric used in CGBayesNets. The BDeu score requires prior knowledge (i.e., equivalent sample size priors) which are typically arbitrarily set to 1; however, multiple studies have shown the sensitivity of BDeu to these parameters [27, 28], as well as the use of improper prior distributions [29]. Alternatively, BIC score does not depend on the prior over the parameters, thus, an ideal approach for scenarios where prior information is not available or difficult to obtain. Next, in order to maximize the full log-likelihood term we implemented a greedy hill-climbing algorithm. We initialize the structure by first connecting each taxa node at the previous time point (for example *T*_1__*_ti_* in Fig. 1e) to the corresponding taxa node at the next time point *(T*_1__*_ti+1_* in Fig. 1e). We call this setting the *baseline* model since it ignores dependencies between taxa’s and only tries to infer taxa levels based on its levels in the previous time points. Next, we added nodes as parents of a specific node via intra or inter edges depending on which valid edge (i.e., no cycles) leads to the largest increase of the log-likelihood function beyond the global penalty incurred by adding the parameters as measured by the BIC^3^ score approximation

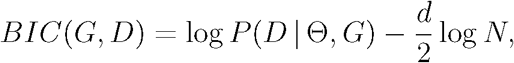

where d = |Θ| is the number of DBN model parameters in *G*, and *N* is the number of time points in *D*. Additionally, we imposed an upper bound limit on the maximum number of possible parents *(maxParents* ∈ {1, 3, 5}) for each bacterial node X (i.e., |**Pa***^G^*(*X*)| ≤ *maxParents)*.

#### Inferring biological relationships

Microbial ecosystems are complex, often displaying a stunning diversity and a wide variety of relationships between community members. These bixological relationships can be broadly divided into two categories: *beneficial*: (including mutualism, commensalism and obligate), or *harmful* (including competition, amensalism and parasitism). Although the longitudinal data sets considered in this study do not provide enough information to further sub-categorize each biological relationship (e.g., mutualism vs. commensalism), we use the learned DBN model from each microbiome data set and inspect each interaction as a means for inferring simple to increasingly complex relationships. For example, consider variable *T*_4__*_ti_* in Fig. 1e. Given that *ti* and *ti*+1 represent the previous time point and the current time point (respectively), the possible inference in this case is as follows: Edges from *T*_4__*_ti_* and *C*_3__*_ti_* (inter edges), and from *T*_2__*_ti_*_+1_ (intra edge) suggest the existence of a temporal relationship in which the abundance of taxa *T*_4_ at a previous time instant and abundance of taxa *T*_2_ at the current time instant, as well as condition *C*_3_ from the previous time instant impact the abundance of *T*_4_ at the current time. We previously stated that *f*(*T*_4__*_ti_*_+1_ | *T*_4__*_ti_, C*_3__*_ti_, T*_2__*_ti_*_+1_) is modeled by *N*(*λ*_0_ + *λ*_1_ × *T*_4__*_ti_* + *λ*_2_ × *C*_3__*_ti_* + *λ*_3_ × *T*_2__*_ti_*_+1_*, σ*_2_). Therefore, inspecting the regression coefficients *λ*_1_*, λ*_2_*, λ*_3_ immediately suggests whether the impact is positive or negative. In this example, the regression coefficients *λ*_1_*, λ*_2_ are positive (*λ*_1_*, λ*_2_ *>* 0) while coefficient *λ*_3_ is negative (*λ*_3_ *<* 0), thus, variables *T*_4__*_ti_* and *C*_3__*_ti_* exhibit positive relationships with microbial taxa *T*_4__*_ti_*_+1_ shown as green edges in Fig. 1e, whereas taxa *T*_2__*_ti_* exhibits a negative interaction with *T*_4__*_ti_*_+1_ shown as a red edge (Fig. 1e). This simple analytic approach enables us to annotate each biological relationship with directional information.

#### Network visualization

All the bootstrap networks^4^ shown are visualized using *Cytoscape* [30] version 3.6.0, using Attribute Circle Layout with Organic Edge Router. An in-house script is used to generate a custom style XML file for each network, encoding the following information in the graph:

- Time *ti* nodes colored in orange
- Time *ti* + 1 nodes colored in blue
- Number of incoming edges directly proportional to node size
- Taxa abundance directly proportional to node transparency
- Clinical and demographic nodes represented with diamond shaped nodes
- Taxa nodes represented with circle shaped nodes
- Solid edges represent inter edges (i.e., from time slice *ti* to *ti* + 1)
- Dashed edges represent intra edges
- Positively weighted edges are colored in green
- Negatively weighted edges are colored in red
- Regression coefficient directly proportional to edge thickness
- Bootstrap value directly proportional to edge transparency

Also the regression coefficients corresponding to edge thickness were normalized as follows: Let *y* be a microbial taxa node with continuous taxa parents *u*_1_, ··· *,u_k_* modeled by

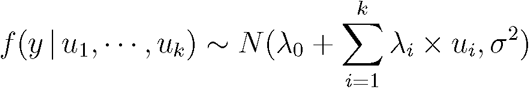

where *λ*_1_, ··· *,λ_k_* are the corresponding regression coefficients for *u*_1_, ··· *,u_k_* as previously described in this section. The normalized regression coefficients 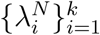 are defined as

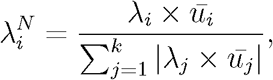

where *ū_i_* is the mean abundance of taxa *u_i_* across all samples.

## 3. Results

Fig. 1 presents the computational pipeline we developed for aligning and learning DBNs for microbiome and clinical data. We start by estimating a cubic spline from the observed abundance profile of each taxa (Fig. 1b). Next, we determine an alignment which allows us to directly compare temporal data across individuals (Fig. 1c), as well as filter out abnormal and noisy samples (Fig. 1d). Finally, we use the aligned data to learn causal dynamic models that provide information about interactions between taxa, their impact, and the impact of clinical variables on taxa levels over time (Fig. 1e-f).

We applied our methods to study longitudinal data sets from three human microbiome niches: infant gut, vagina and oral cavity (see Methods for full descriptions). In addition to the differences in the taxa they profile, these data sets vary in the number of subjects profiled (ranging from 18 to 58, in the number of time points they collected, the overall number of samples and time series that were studied, etc. Thus, they provide a good set to test the generality of our methods and their usefulness in different microbiome studies.

### Infant gut alignments captures gestational age at birth

Below, we discuss in detail the improved accuracy of the learned dynamic models due to use of *temporal alignments*. However, even before using them for our models, we wanted to test whether the alignment results agree with biological knowledge. For this, we used the infant gut data. Infant gut microbiota goes through a patterned shift in dominance between three bacterial populations (*Bacilli* to *Gammaproteobacteria* to *Clostridia*) in the weeks immediately following birth. La Rosa *et al*. [5] reported that the rate of change is dependent on maturation of the infant highlighting the importance of post-conceptional age as opposed to day of life when analyzing bacterial composition dynamics in preterm infants. We found that our alignment method is able to capture this rate of change without explicitly using gestational or post-conceptional age.

Fig. 2 shows the relationship between alignment parameters *a* and *b* (from the transformation function 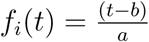 described in Methods) and the gestational age at birth for each infant in the gut microbiome data set. Each aligned infant sample is represented by a blue circle where the x-axis shows 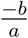 and y-axis shows the gestational age at birth. As can be seen, the alignment parameters are reasonably well correlated with gestational age at birth (Pearson’s correlation coefficient = 0.35) indicating that this method can indeed be used to infer differences in rates between individuals.

**Figure 2:**
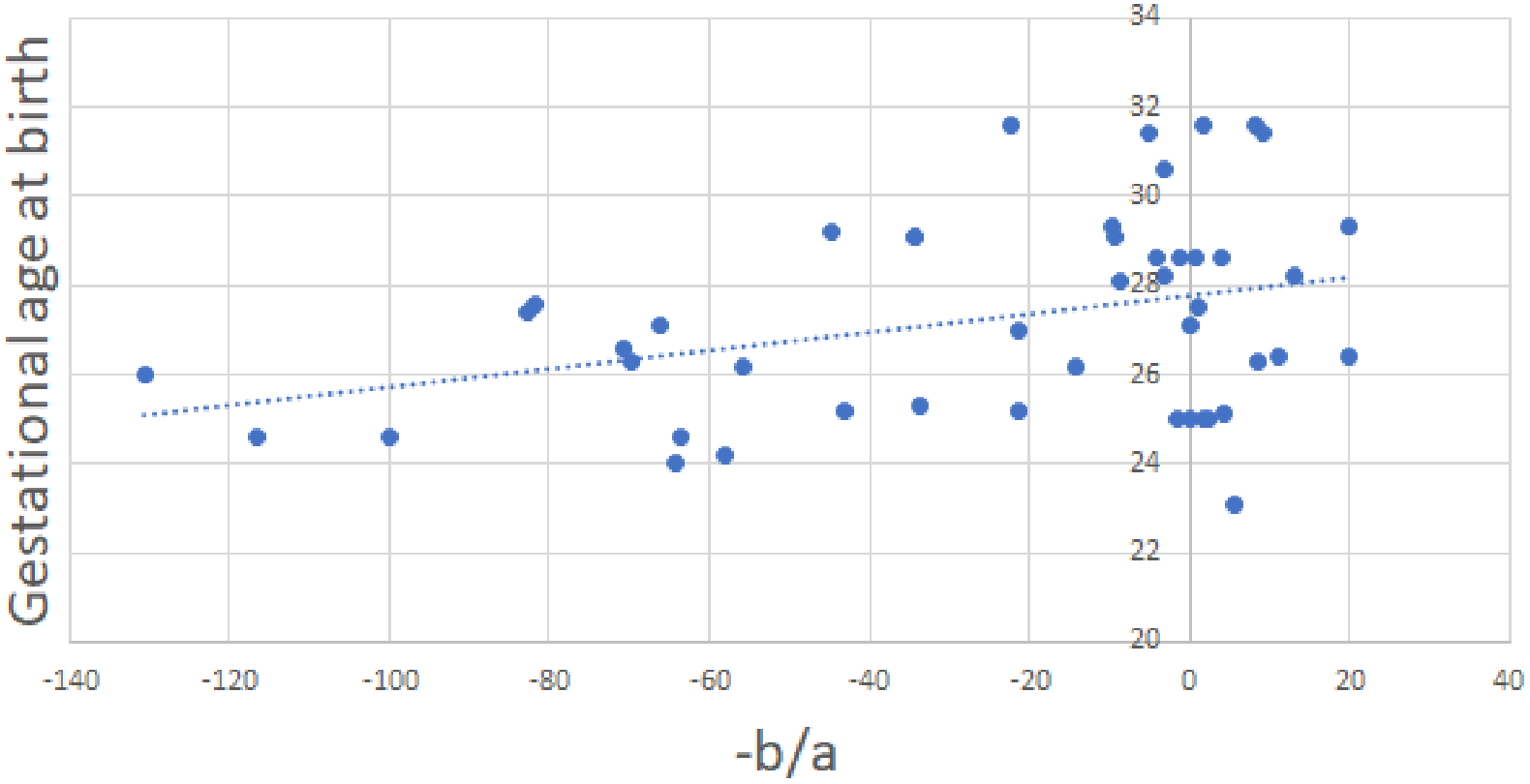
Relationship between alignment parameters and gestational age at birth. Figure shows the relationship between alignment parameters *a* and *b* and gestational age at birth for the aligned infant gut microbiome data set. Each blue dot represent an aligned infant sample *i* where x-axis shows 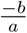 from transformation function 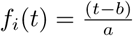 and y-axis shows the gestational age at birth of infant *i*. Pearson correlation coefficient = 0.35.

### Resulting dynamic Bayesian network models

We next applied the full pipeline to learn DBNs from the three microbiome data sets under study. In particular, we use longitudinal data sets from three human microbiome niches: infant gut, vaginal and oral cavity as described in Methods. In this section, we highlight the overall characteristics of the learned DBN for each aligned and filtered microbiome data set (Figure 3 and Additional file 5: Figure S3). In these figures the nodes represent taxa and clinical (or demographic) variables and the directed edges represent temporal relationships between them. Several triangles were also observed in the networks. In some of the triangles, directed edges to a given node were linked from both time slices of another variable. We will refer to these as *directed triangles*.

**Figure 3:**
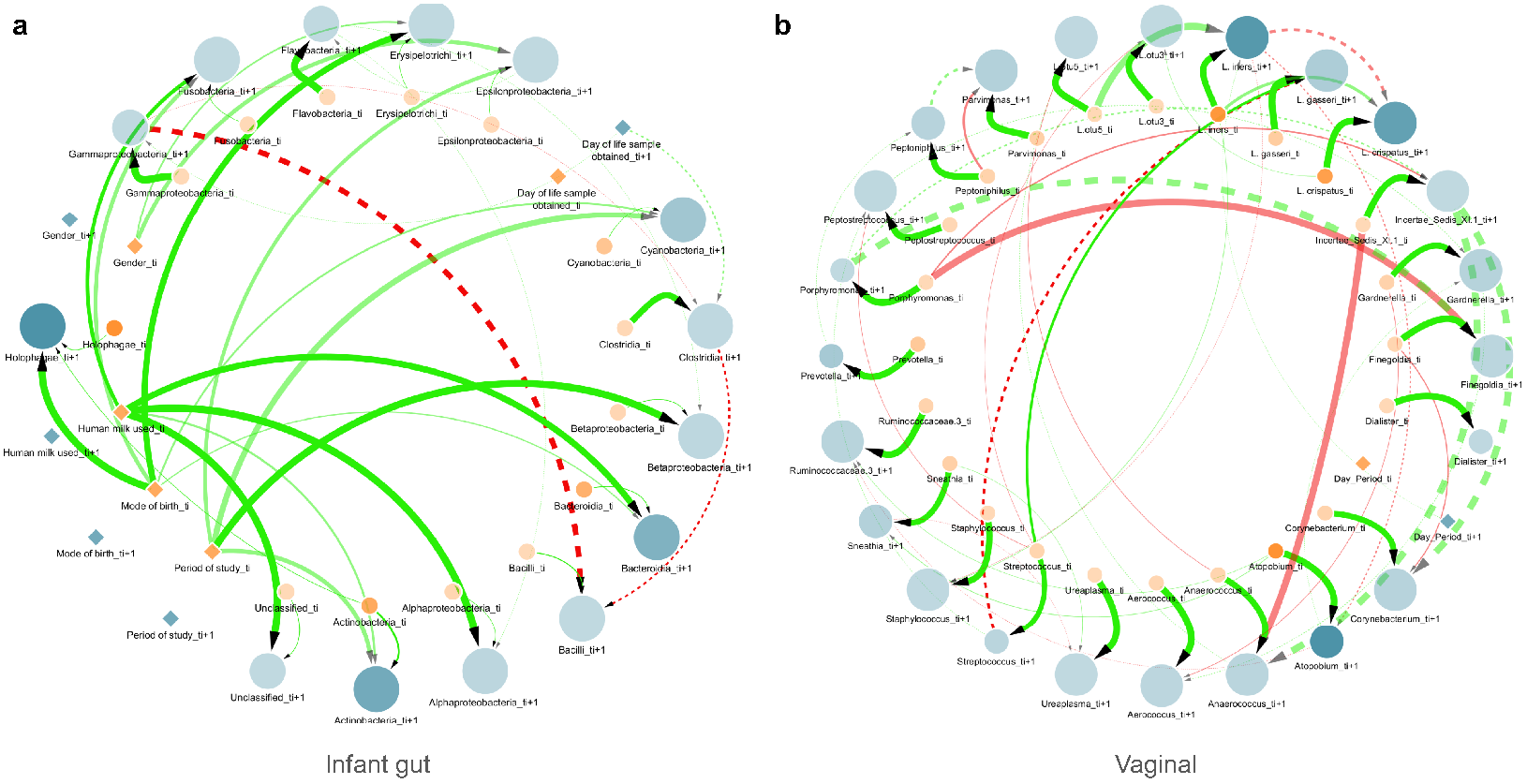
Dynamic Bayesian network for two representative data sets. Figure shows two consecutive time slices *ti* (orange) and *ti* + 1 (blue), where nodes are either microbial taxa (circles) or clinical factors (diamonds). Nodes size is proportional to in-degree whereas taxa nodes transparency indicates mean abundance. Additionally, dotted lines denote *intra edges* (i.e., directed links between nodes in same time slice) whereas solid lines *denote inter edges* (i.e., directed links between nodes in different time slices). Edge color indicates positive (green) or negative (red) temporal influence and edge transparency indicates strength of bootstrap support. Edge thickness indicates statistical influence of regression coefficient as described in Network visualization**. a** — Learned DBN for the aligned infant gut microbiome data at a sampling rate of 3 days and maxParents = 3. **b** — Learned DBN for the aligned vaginal microbiome data at a sampling rate of 3 days and *maxParents* = 3.

#### Infant gut

The learned DBN model for the infant gut microbiota data set at a sampling rate of 3 days and *maxParents* = 3 was computed. It contains 19 nodes per time slice (14 microbial taxa, 4 clinical and 1 demographic variable nodes) and 39 directed edges (31 inter edges and 8 intra edges) with no directed triangles as shown in Fig. 3a. Since we only learn temporal conditional dependence (i.e., incoming edges) for taxa nodes at time slice *i* + 1, the maximum number of possible edges is 14 × *maxParents* = 42, thus, most of the taxa nodes (11 out of 14) have reached the maximum number of parents allowed (i.e., *maxParents* = 3). Additionally, the majority of these temporal relationships are between microbial taxa. In particular, the model includes several interactions between the key colonizers of the premature infant gut: *Bacilli*, *Clostridia* and *Gammaproteobacteria*. Furthermore, the only negative interactions learned by the model comprise these microbes which are directly involved in the progression of the infant gut microbiota. Also, the nodes for gestational age at birth and post-conceptional age at birth are not shown because they are isolated from the rest of the network, without any single edge. Overall, these trends strongly suggest that the DBN is capturing biologically relevant interactions between taxa.

#### Vaginal

As with the gut microbiome data set, we learned a DBN model for the vaginal microbiome data at a sampling rate of 3 days and *maxParents* = 3 (Fig. 3b). The resulting DBN is comprised of 24 nodes per time instance (23 taxa and 1 clinical) and 58 edges (40 inter edges and 18 intra edges). Additionally, 12 directed triangles involving taxa nodes were observed. In preliminary analyses, additional clinical and demographic attributes (e.g., Nugent category, race and age group) resulted in networks with these variables connected to all taxa nodes, thus, removed from further analysis. Specifically, we estimated the degree of overfitting of these variables by learning and testing DBN models with and without them. This resulted in the DBN shown in Fig. 3b which exhibited lowest generalization error. In this case, the maximum number of potential edges between bacterial nodes is 24 × *maxParents* = 72; however, only 16 out of 24 taxa nodes reached the threshold on the maximum number of parents. Among all the 58 edges, only one interaction *Day_Period_ti+1* to *L. iners_ti+1* involves a clinical node whereas the remaining 57 edges (including 15 negative interactions) captured temporal relationships among microbial taxa. This mixture of positive and negative interactions between taxa provides evidence of the DBNs ability to capture the complex relationships and temporal dynamics of the vaginal microbiota.

#### Oral cavity

We learned a DBN with the longitudinal tooth/gum microbiome data set with a sampling rate of 7 days and *maxParents* = 3. Additional file 5: Figure S3 shows the learned DBN which contains 20 nodes for each time slice (19 taxa and 1 clinical) and 52 edges (33 inter edges and 19 intra edges) out of 57 possible edges. In addition 2 directed triangles were observed involving taxa nodes. Here, the DBN model includes multiple positive and negative interactions among early colonizers (e.g., *Veillonella* and *H. parainfluenzae*) and late colonizers (e.g., *Porphyromonas*) of the oral microbiota which are supported by previous experimental studies [31].

### Comparisons to prior methods

To evaluate the accuracy of our pipeline and to compare them to models reconstructed by prior methods published in the literature [32, 10], we used a per-subject cross-validation with the goal of predicting microbial taxon abundances using the learned models. In each iteration, the longitudinal microbial abundance profile of a single subject was selected as the test set, and the remaining profiles were used for building the network and learning model parameters. Next, starting from the second time point, we used the learned model to predict an abundance value for every taxa in the test set at each time point using the previous and current time points. Predicted values were normalized to represent relative abundance of each taxa across the microbial community of interest. Finally, we measured the average predictive accuracy by computing the mean absolute error (MAE) for the selected taxon in the network. We repeated this process (learning the models and predicting based on them) for several different sampling rates, which ranged from 1 up to 28 days depending on the data set. The original and predicted microbial abundance profiles can be compared as shown in Fig. 1f. The average MAE for predictions on the three data sets are summarized in Fig. 4(a-c). For each data set, error plots are shown for ten different methods. Along with two of our DBNs (one with and one without alignments), four methods with and four without alignments were compared. These are further described below.

**Figure 4:**
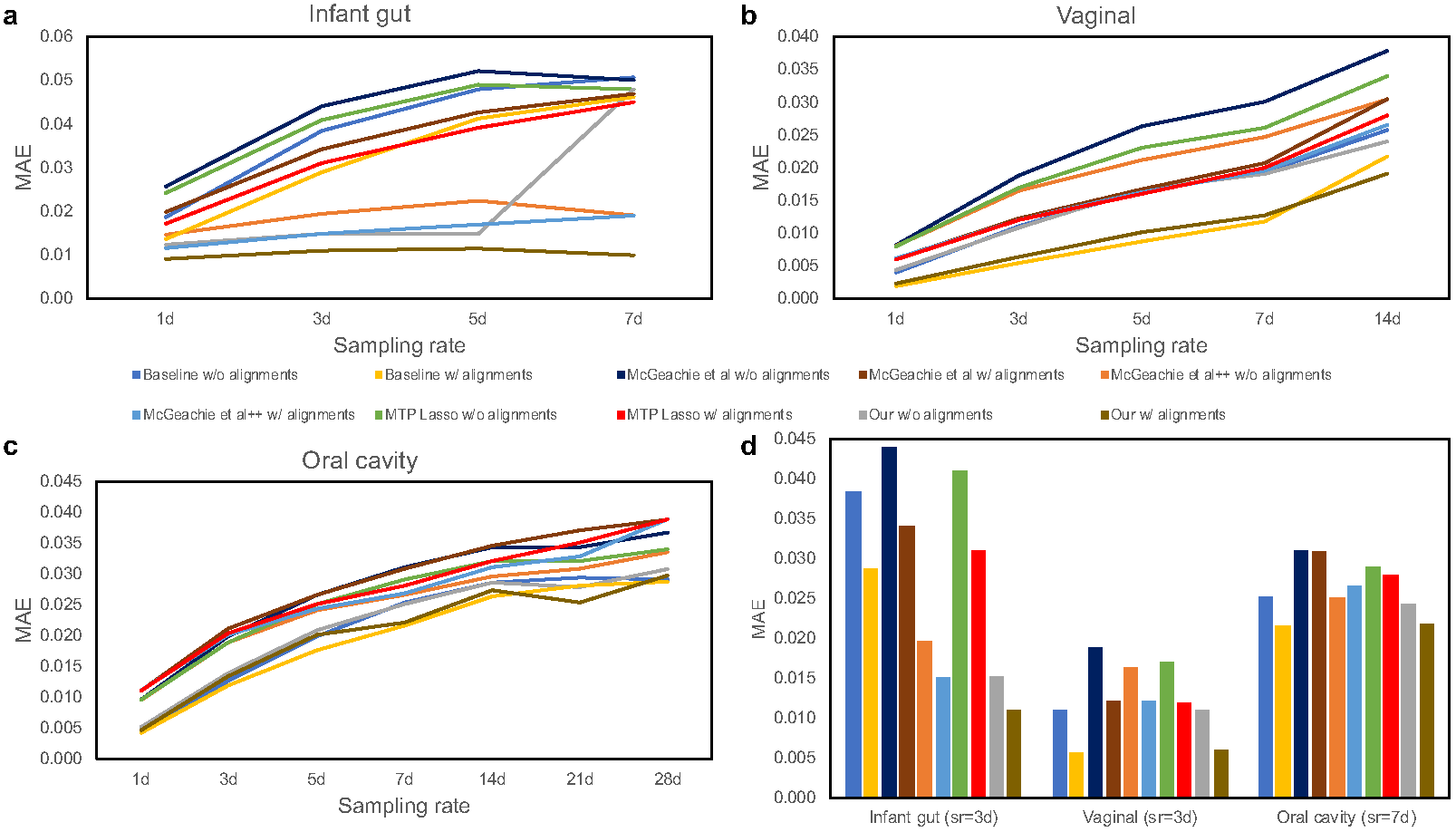
Comparison of average predictive accuracy between methods on the filtered data sets. Figure shows the average MAE of our proposed DBN models against a baseline method and previously published approaches as a function of sampling rates where *d* denotes day(s). Additionally, each method is run on the unaligned and aligned data sets. **a** — Performance results for infant gut microbiome data. **b** — Performance results for vaginal microbiome data. **c** — Performance results for oral cavity microbiome data. **d** — Performance results for each data set for a sampling rate (*sr*) that most closely resembles the originally measured time points.

First, we compared the DBN strategy to a naive (baseline) approach. This baseline approach makes the trivial prediction that the abundance value for each taxa *A* at any given point is exactly equal to the abundance measured at the previous time point. Given that measured abundances are continuous variables, this turns out to be an extremely competitive method and performs better than most prior methods for the data sets we tested on. Next, we compared our DBNs to three other methods suggested for modeling interactions among taxa: (a) McGeachie *et al*. [10] developed a different DBN model where network learning is estimated from the BDeu scoring metric [23] (instead of MLE), (b) McGeachie *et al*.++ an in-house implementation that extends McGeachie *et al*.’s method to allow for intra edges during structure learning, and (c) MTPLasso [32] that models time-series microbial data using a gLV model. In all cases, we used the default parameters as provided in the original publications.

As can be seen, our method outperforms the baseline and previous methods for the infant gut data. It also performs favorably when compared to baseline on the other two data sets. Temporal alignments improved the predictive performance over unaligned samples across gut and vaginal microbiomes by about 1–4 percentage points. In particular, a two-tailed t-test indicates significant (denoted by *) performance improvements for most sampling rates (*infant gut*: *p* − *value* =0.043* for 1*d*, *p* − *value* =0.034* for 3*d*, *p* − *value* =0.109 for 5*d*, and *p* − *value <* 1.00*E* − 05* for 7*d*; *vaginal*: *p* − *value <* 1.00*E* − 06* for 1*d*, *p* − *value <* 1.00*E* − 05* for 3*d*, *p* − *value* =5.50*E* − 05* for 5*d*, *p* − *value* =3.10*E* − 03* for 7*d*, and *p* − *value* =0.097 for 14*d*). On the other hand, alignments did not show significant predictive performance improvements on the oral data set and is consistent with previous analysis on the same data set [12]. Surprisingly, the simple baseline approach outperforms all previously published methods: McGeachie *et al*. [10] and MTPLasso [32] across the three data sets. Finally, Fig. 4d highlights the MAE results for a sampling rate that most closely resembles the originally measured time points.

### Anomaly detection using alignment

When analyzing large cohorts of microbiome data, it is important to implement a strategy to remove outliers as these can affect our ability to generalize from the collected data. As discussed in Methods, we can use our alignment error *E_M_* score to identify such subjects and remove them prior to modeling. In the context of the gut data set, this resulted in the identification of two infant samples: Subjects 5 and 55 (highlighted in red within Additional file 6: Figure S4a) which are likely processing errors, contaminated samples, or just natural anomalies. Sample 55 has been previously identified as a likely abruption event by McGeachie *et al*. [10] using a different approach. Similarly, Additional file 6: Figure S4b shows the distribution of alignment errors *E_M_* for the vaginal microbiome data. In this case, we remove 6 sub-samples from 4 different women (highlighted in red). We note that there were no outliers identified in the oral cavity microbiome data set. When learning DBNs following the filtering we obtain even better models. Additional file 7: Figure S5 compares the average MAE results of our proposed DBN model between the unfiltered and filtered samples for the gut and vaginal data sets. As can be seen, a large performance improvement is observed for the gut data while a slight improvement is observed for the vaginal data when removing the outliers. These results suggest that even though the method uses less data to learn the models, the models that it does learn are more accurate.

## 4. Discussion

### The power of temporal alignments

We developed a pipeline for the analysis of longitudinal microbiome data and applied it to three data sets profiling different human body parts. To evaluate the reconstructed networks we used them to predict changes in taxa abundance over time. Interestingly, ours is the first method to improve upon a naive baseline (Fig. 4). While this does not fully validate the accuracy of the models, it does mean that the additional interactions determined by our method contribute to the ability to infer future changes and so at least some are likely true.

As part of our pipeline we perform temporal alignment. While ground truth for alignments is usually hard to determine, in one of the data sets we analyzed we could compare the alignment results to external information to test its usefulness. In the context of the infant gut data, it has been shown that using day of life as the independent variable hinders the identification of associations between bacterial composition and day of sampling. Therefore, previous work have re-analyzed the premature gut microbiota with post-conceptional age, uncovering biologically relevant relationships [5]. By using alignment we were able to correct for this difference without the need to rely on the external age information. In addition to the results presented in Fig. 2, the learned DBN in Fig. 3a does not show any relationships to post-conceptional age or gestational age at birth indicating that our alignment was able to successfully compensate for. While for this data such correction could have been made using post-conceptional age, in other cases the reason for the rate change may not be obvious and without alignment it would be hard to account for such hidden effects.

### Uncovering biological relationships

We next discuss in more detail the learned DBN models.

#### Infant gut

As mentioned in Results, the only negative relationships identified supports the known colonization order, that is, a shift in dominance from *Bacilli* to *Gammaproteobacteria* to *Clostridia*) [5], as the infant goes through the first several weeks of life. These edges show incoming negative relationships to *Bacilli* from *Gammaproteobacteria* and *Clostridia*. In particular, an increase in the abundance of the parents is associated with a decrease in the abundance of the child. The negative edge from *Gammaproteobacteria* to *Clostridia* agrees with previous findings where *Clostridia*’s abundance is found to increase at a gradual rate until it peaks at post-conceptional age between 33 and 36 weeks whereas *Gammaproteobacteria* decreases as infants age [5, 10]. This relationship is also confirmed by the edges from *Day of life* to *Gammaproteobacteria* and *Clostridia* (Fig. 3b). Moreover, the DBN model indicates a relationship between breastfeeding and *Actinobacteria*, *Bacteroidia*, and *Alphaproteobacteria*. These bacteria are known to be present in breast milk which is known to heavily influence and shape the infant gut microbiome [33].

#### Vaginal

It has been established that microbial composition can change dramatically during the menses cycle and later return to a ‘stable’ state before the next menstrual period [34, 35]. Previous studies have identified a subset of individuals in this data set as exhibiting a microbial composition dominated by *L. crispatus* with a notable increase of *L. iners* around the start of each menstrual period [4, 34] (Additional file 2: Figure S1a). These interactions were also captured by the learned DBN model in the form of a directed triangle involving *L. crispatus* and *L. iners* (Fig. 3b). On the other hand, subjects from another group were characterized as dominated by *L. gasseri* coupled with shifts to *Streptococcus* during menstruation [4]. These relationships were also captured by the DBN. The edge from the *Day Period* to *L. iners* strengthens this relationship. Furthermore, while *L. iners* has a lower protective value than the other *Lactobacillus* [36], the negative edge between *L. iners* and *Atopobium* suggests a relationship related to environment protection. Also, the positive edge from *Atopobium* to *Gardnerella* is supported by the synergy observed between these two taxa in bacterial vaginosis [37]. Finally, it is important to note that the shifts and composition of the vaginal microbiome vary considerably between each subject.

#### Oral

For oral microbiomes, several *Streptococcus* species, including *S. oralis*, *S. mitis*, *S. gordonii*, and *S. sanguis* are well known as early colonizers lying close to the tooth pellicle [31]. While our DBN cannot identify specific species, it suggests interactions between some species of *Streptococcus* and other later colonizers in the oral microbiome such as *Porphyromonas* and *Prevotella*. The DBN also provided novel predictions, for example taxa *Granulicatella* is interacting with *Veilonella*.

### Triangles in DBNs

An interesting aspect shared by all of the DBNs discussed above is the fact that they contain triangles or feed-forward loops. In particular many of these directed triangles are created from nodes representing both time slices of another variable, but with different signs (one positive and the other negative). For example, microbial taxa *L. crispatus* displays a directed triangle with another taxa *L. iners* in the vaginal DBN (Fig. 3b). In this triangle, positive edges from *L. iners_ti* interact with *L. iners_ti+1* and *L. crispatus_ti+1* whereas a negative edge connects *L. iners ti+1* to *L. crispatus ti+1*.

The triangles in the DBNs represent a relationship where the abundance of a child node cannot be solely determined from the abundance of a parent at one time slice. Instead, information from both the previous and the current time slices is needed. This can be interpreted as implying that the child node is associated with the *change* of the abundance values of the parents rather than with the absolute values which each node represents.

### Limitation and future work

While our pipeline of alignment followed by DBN learning successfully reconstructed models for the data sets we looked at, it is important to understand the limitation of the approach. First, given the complexity of aligning a large number of individuals, our alignment method is based on a greedy algorithm, thus, it is not guaranteed to obtain the optimal result. Even if the alignment procedure is successful, the DBN may not be able to reflect the correct interactions between taxa. Issues related to sampling rates can impact the accuracy of the DBN (missing important intermediate interactions) while on the other hand if not enough data is available the model can overfit and predict non-existent interactions.

Given these limitations we would attempt to improve the alignment method and its guarantees in future work. We are also interested in studying the ability of our procedure to integrate additional molecular longitudinal information including gene expression and metabolomics data which some studies are now collecting in addition to the taxa abundance data [38]. We believe that our approach for integrating information across individual in order to learn dynamic models would be useful for several ongoing and future studies.

## 5. Conclusions

In this paper, we propose a novel approach to the analysis of longitudinal microbiome data sets using dynamic Bayesian networks with the goal of eliciting temporal relationships between various taxonomic entities and other clinical factors describing the microbiome. The novelty of our approach lies in the use of temporal alignments to normalize the differences in pace of biological processes inherent within different subjects. Additionally, the alignment algorithm can be used to filter out abruption events or noisy samples. Our results show that microbiome alignments improve predictive performance over previous methods and enhance our ability to infer known and potentially novel biological and environmental relationships between the various entities of a microbiome and the other clinical and demographic factors that describe the microbiome.

## 7. Acknowledgements

The authors would like to thank Jun Ding for his assistance in implementing the temporal alignment algorithm.

## Footnotes

3 We also computed AIC score (i.e., *AIC*(*G, D*) = log *P*(*D*|Θ;*G*) − *d*) but it was consistently outperformed by BIC score.

4 For each data set, we ran 500 bootstrap realizations and only reported edges with bootstrap support of at least 50% in the consensus DBN.

